# Lrrc55 modulates beta-cell resilience and calcium transporter expression under metabolic and physiologic stress

**DOI:** 10.1101/2025.11.03.686179

**Authors:** Marle Pretorius, Raneet Kahlon, Qianyu Xie, Carol Huang

## Abstract

During pregnancy, pancreatic β-cell mass and function are increased in adaptation to insulin resistance; this requires the action of prolactin receptor signaling. Recently, we discovered the Lrrc55, an auxiliary subunit of the voltage- and calcium-activated potassium channel (BK channel), is a prosurvival factor in β-cell, and its expression and highly and specifically up regulated in the pancreatic islets during pregnancy. We found that overexpression of Lrrc55 protects β-cells from glucolipotoxicity (GLT)-induced apoptosis. The protective effect of Lrrc55 is associated with dampening of the ER stress response and preservation of the releasable pool of calcium in the ER. Thus, we hypothesized that Lrrc55 protects β-cells from GLT-induced ER stress and apoptosis by regulating ER calcium handling. Here, we report that Lrrc55 restored the GLT-mediated decrease in expression levels the insulin regulators, *Pdx-1* and *MafA*, as well as the ER calcium regulator SERCA. Lrrc55 also attenuated the GLT-induced increase in expression of the ER calcium channel RyR2. Lrrc55 also attenuated GLT-mediated increase in protein expression of pro-apoptotic molecules CHOP and IRE1α, and increased activation of the anti-apoptotic molecule Akt. However, Lrrc55 does not alter the activity levels of SERCA or IP3R under physiologic conditions. Transgenic mice with global deletion of Lrrc55 (Lrrc55^−/−^) are more susceptible to streptozotocin-induced diabetes. Surprisingly, Lrrc55^−/−^ mice are more glucose tolerant and secreted more insulin during pregnancy. Together, these results suggest a role for Lrrc55 in maintaining expression of the ER calcium regulators in the presence of GLT but it may negatively regulate insulin secretion.

## Introduction

Pancreatic islets adapt to metabolic stressors such as diet-induced obesity and pregnancy by up regulating beta-cell proliferation, insulin synthesis, and secretion. Previously, we found that during pregnancy, placental lactogens and prolactin act through the prolactin receptor (PRLR) to increase β-cell mass and increase insulin release. Mice with either global or β-cell specific PRLR deletion experienced impaired glucose tolerance during pregnancy, mainly due to a defect in the adaptive upregulation of β-cell number^1,2^. This defect was accompanied by downregulation of signaling molecules known to regulate β-cell proliferation, namely Menin, IRS-2, p21 and Akt^3^. Through microarray analysis, *Lrrc55* emerged as a potential new target of PRLR. Lrrc55 is an auxiliary subunit of the BK channel^4,5^, and is up regulated by >60-fold in islets during pregnancy^6^. This pregnancy-associated upregulation is specific to Lrrc55, as the expression of other auxiliary subunits of the BK channel, namely Lrrc26 and Lrrc38, are not altered during pregnancy. Overexpression of Lrrc55 in β-cell protected them from fatty-acid-induced cellular dysfunction and apoptosis. This was associated with upregulation of pro-survival genes and down regulation of pro-apoptosis genes, with attenuation of the endoplasmic reticulum (ER) stress pathway. Overexpression of Lrrc55 also preserved insulin content in fatty-acid-treated islets. One interesting observation was that Lrrc55 protected β-cells from fatty-acid-induced calcium depletion from the ER stores^6^. As ER calcium depletion can initiate ER stress and apoptosis, delineating how Lrrc55 modulates ER calcium homeostasis will provide insight into its function as a pro-survival factor.

ER calcium concentration is regulated by sarco/endoplasmic reticulum Ca^2+^ - ATPase (SERCA), which actively transports calcium from the cytosol into the ER, as well as two receptor-type membrane proteins, the ryanodine receptor (RyR) and the Inositol Triphosphate Receptor (IP3R)^7-10^. Inhibition of SERCA has been shown to cause ER stress, and interestingly, this effect was attenuated by blocking IP3R and RyR^11^. Since we found that Lrrc55-mediated pro-survival function coincide with preservation of ER calcium stores^6^, the goal of this study was to explore how Lrrc55 affects the expression and activity of calcium transporters, and to assess its impact on glucose homeostasis in vivo.

## Methods

### Ethical approval

All experimental procedures were approved by the Animal Use Review Committee at the University of Calgary, in compliance with the standards of the Canadian Council on Animal Care and the ARRIVE guidelines.

### Mice

The global Lrrc55-knockout C57Bl/6 mice were a generous gift from Dr. Jiusheng Yan of the M. D. Anderson Cancer Center at the University of Texas. Mice were maintained on a 12-hour light and 12-hour dark cycle with no restrictions to water or food. Heterozygous male and female mice were interbred to generate homozygous experimental mice. Mice were studied at 3-6 months of age. For timed pregnancy experiments, the day a vaginal plug is detected is designated as gestational day 1. To induce diabetes during pregnancy, streptozotocin (100mg/kg/d) was given on days 13 and 14 of pregnancy, and non-fasting blood glucose was measured twice weekly, starting 7 days after the 2^nd^ dose of streptozotocin. Non-fasting blood glucose > 11 mM for 2 consecutive weeks were defined as having diabetes.

### Materials

All chemical reagents were purchased from Sigma Millipore Canada, unless otherwise specified. Cell culture reagents were purchased from Thermo Fisher Scientific.

### Physiological testing for glucose homeostasis

Glucose tolerance test (GTT)^1^: Mice were fasted overnight and at 8am, glucose solution was given by oral gavage for an oral glucose tolerance test (OGTT, 2g/kg body weight 40% D-glucose in 0.9% normal saline). Blood glucose was sampled from the tail vein by a glucose meter (One Touch Verio IQ) at times 0, 10, 15, 30, 45, 60, and 120 minutes after glucose administration. Serum (∼30 μl) was obtained at times 0, 10, and 30 minutes after glucose administration for insulin measurement by ELISA (ALPCO).

Insulin tolerance test (ITT)^1^: Mice were fasted for 4 hours, then insulin (0.5 units/kg body weight, NovoRapid) was injected intraperitoneally and blood glucose was sampled from the tail vein at times 0, 15, 30, 45, and 60 minutes after insulin injection.

### Islet isolation

Pancreatic islets were isolated from non-fasted mice as previously described^6^. Pancreas was distended using 2.5 ml/pancreas collagenase Type XI (0.5mg/mL (Roche: Catalog #C7657) in Hank’s balanced salt solution (HBSS)), removed by blunt dissection, and cleared of fat. The distended pancreas was then incubated at 37°C for 13-15 minutes under constant agitation in a 50mL falcon tube. Islets were dislodged from the exocrine pancreas by vigorous tapping of the tube by hand for ∼10-15 seconds. The digestion is stopped by adding 35 mL of ice cold HBSS (with 1mM calcium), and let the islets settle to the bottom of the tube by gravity, on ice, for ∼ 7 minutes. 20mL of the supernatant was then carefully removed, then topped up with 20 mL of fresh, ice cold HBSS. Islets were then hand-picked and 30 islets were cultured overnight in RPMI 1640 medium with glutamine (Hyclone, catalog number SH3002701), supplemented with 10% fetal bovine serum (Gibco, catalog number 12483020) and 1U/100mL of penicillin-streptomycin (Gibco, catalog number 15070063) at 37°C and 5% CO_2_ for in vitro glucose-stimulated insulin secretion assay the next day. The remaining islets were flash-frozen and stored in −80°C for RNA or protein purification.

### Islet culture and adenovirus infection

Hand-picked islets were cultured in RPMI 1640 supplemented with 10% FBS and 1U/mL penicillin/streptomycin. For islet infection with Adenovirus, whole islets were gently dissociated by incubation for 5–7 min in EDTA (2 mM) at 37°C. Islets were then infected with a 100:1 multiplicity of infection (MOI) of Ad-GFP or Ad-Lrrc55 in growth media. They were cultured in virus-containing media for a total infection time of 48 hours, with a top-up of fresh growth media after 24 hours of infection^6^.

### RT-qPCR

Sample cDNA was synthesized from 1µg RNA using the Quantitect Reverse Transcription Kit for SYBR green quantitative PCR (Qiagen), then diluted 1:5 in water. RT-qPCR reactions were carried out in duplicate using the Quantitect SYBR Green PCR kit (Qiagen) and the QuantStudio 6 Flex Real-Time PCR System (Applied Biosystems) with annealing temperature of 61°C. Relative expression levels were normalized against housekeeping gene PGK1 due to its low variability across samples of islets from pregnant and non-pregnant mice^12^. Each primer pair was validated by ensuring that melt curves contained a single peak and that amplified products contained a single gene product of the predicted amplicon length. The 2−ΔΔCt method was used to analyze relative fold-changes in gene expression.

### Cell Culture

HEK 293T cells (from ATCC, CRL-3216) were used for the SERCA assay. Cells were seeded at 2.5×10^5^ cells/10-cm plate in DMEM media (11mM glucose, 110mg/L Sodium Pyruvate) supplemented with 1U/mL Penicillin-Streptomycin and 10% FBS, cultured at 37°C and 5%CO_2_. Flp-In T-REx-293T cells overexpressing IP3R (generously given by the Chen Laboratory, University of Calgary) were used for the IP3R assay. Cells were seeded at 5×10^5^ cells/plate in DMEM media (11mM glucose, 110mg/L Sodium Pyruvate) supplemented with 1U/mL Penicillin-Streptomycin and 10% FBS, 200mM L-Glutamine, and 1X MEM Non-Essential Amino Acids, cultured at 37°C and 5% CO_2_.

### Transfection

HEK 293T cells or Flp-In T-REx-293T cells overexpressing were transfected 24 hours after plating, using the calcium-phosphate precipitation method. For IP3R activity assays, 4µg of DNA per plasmid was transfected per plate. For SERCA activity assays, 2µg of DNA per plasmid was transfected per plate. DNA was diluted into 200µL per plate 125µM CaCl_2_, then added to 200µL per plate of 2X HEPES Solution (pH 7.04, 274mM NaCl, 1.8mM Na_2_HPO_4_, and 50mM HEPES). 320µL per plate of this mixture was added dropwise directly to the cells in media and cells were incubated for 48 hours. For IP3R activity assays, 4µg of DNA per plasmid was transfected per plate. For SERCA activity assays, 2µg of DNA per plasmid was transfected per plate.

### SERCA Activity Assay

G-CEPIA1er, a genetically encoded ER calcium indicator was used in a single cell assay to test SERCA activity, in collaboration with the Jonathan Lytton Laboratory at the University of Calgary, based on the assay developed and outlined by Bovo et al. ^13^. In this assay, HEK 293T cells were transfected (24 hours after seeding at about 20-40% confluence) with 2µg each of plasmids containing G-CEPIA1er, SERCA2b, RyR2, and either Lrrc55 or an empty pcDNA3.1 vector as a negative control. Twenty-four hours after transfection, cells were detached by gentle pipetting and transferred to imaging coverslips (15mm round, coated with Poly-L-Lysine). After an additional 24 hours, G-CEPIA-1 signal was measured through confocal microscopy perfusion with various solutions (detailed below) to alter calcium signalling.

Coverslips were mounted onto an inverted microscope in an Extra-Cellular Matrix (ECM) Buffer (pH 7.4, 135mM NaCl, 5mM KCl, 1mM MgCl2, 25mM HEPES, 1mM CaCl2, and 10mM D-glucose) during confocal optimization. Cells were then permeabilized by perfusion with 0.05% Saponin in an Intracellular Matrix (ICM) Buffer (pH 7.4, 125mM, 3mM MgCl_2_, 25mM MOPS, 3mM ATP, 0.5mM EGTA, and 0.45mM CaCl_2_). Cells were perfused with ICM Buffer containing zero calcium (ICM-zero) for 3 minutes to deplete extracellular calcium, then 5mM caffeine in ICM-zero for 2 minutes to deplete ER calcium to estimate the minimum ER calcium store (F_min_). Cells were then perfused with 15uM Ruthenium Red + 1mM Tetracaine in ICM Buffer (RyR Inhibition Cocktail) for 5 minutes to inhibit the RyR. The final profusion was with 5mM CaCl_2_ and 5µM of Ionomycin in ICM Buffer for 2 minutes. This step maximally increases both cytosolic and ER calcium such that a maximum ER calcium store level (F_max_) can be measured. The G-CEPIA1er fluorescence was taken from the time of signal increase due to perfusion with RyR Inhibition Cocktail, to the time of maximal signal (F_max_). The slope of this signal increase was used as a measure of SERCA activity for each single cell trace.

### IP3R Activity Assay

G-CEPIA1er was used in a cell suspension assay to test IP3R activity, in collaboration with the Chen Laboratory (University of Calgary). Flp-In T-REx-293T cells overexpressing IP3R were transfected with plasmids containing G-CEPIA1er, SERCA2B, RyR2, and either FLAG-Lrrc55 or a negative control vector (pcDNA(-). SERCA2B was the SERCA variant chosen due to its selective expression in pancreatic β-cells^14 15^. RyR2 was introduced into these cells to enable complete and reversible emptying of the ER calcium stores by caffeine ^13^.

After 24 hours of transfection, IP3R overexpression was induced by changing growth media to induction media (1 μg/ml tetracycline, Sigma). After 22 hours of induction, cells were washed with PBS, then removed from plates by gentle pipetting and pelleted by centrifugation at 1,000 rpm for 2 mins (Beckman TH-4 rotor). The cells were permeabilized with 50 μg/ml saponin in a complete intracellular-like medium (ICM, 125 mM KCl, 19 mM NaCl, and 10 mM HEPES, 2 mM ATP, 2 mM MgCl2, 0.05 mM EGTA, and 200 nM free Ca2+, pH7.4 with KOH) at room temperature (23°C) for 6 mins. Cell pellets were collected and washed with ICM 3 times, then added to 2 ml (final volume) in a cuvette. The fluorescence intensity of G-CEPIA1er at 527 nm was measured (494-nm excitation) before and after injection with 1–1000 nM of IP3 (25 °C, SLM-Aminco series 2 luminescence spectrometer, SLM Instruments). The maximum calcium release was determined by the addition of 1 µM of IP3 plus 4 μM of cyclopiazonic acid. The varying doses of IP3 cause a release of ER calcium within the cells in suspension and the resulting drop in G-CEPIA1er signal is used to approximate IP3R activity.

## Results

### Lrrc55 partially reversed glucolipotoxicity-induced downregulation of genes associated with beta-cell function

Previous work in INS-1 cells^6^ showed that overexpression of Lrrc55 increased insulin content in INS-1 cells. To investigate the mechanism underlying this observation, we measured the expression of Pdx1 and MafA – two genes that regulate insulin gene expression^16-18^. Islets were infected with Ad-GFP (control virus) or Ad-Lrrc55 and exposed to glucolipotoxicity (GLT), induced by exposure to 0.5 mM of palmitate in the presence of 33mM glucose (PAHG) for 72 hrs. This condition was chosen to induce beta-cell dysfunction^6,19,20^, and to mimic the elevated free-fatty-acid and glucose conditions observed in type 2 diabetes. We observed a significant reduction in Pdx-1 and MafA expression in Ad-GFP-infected islets after 72 hours of palmitate exposure, which was partially restored by overexpression of Lrrc55 (Figure 1 A, B).

**Figure 1.**
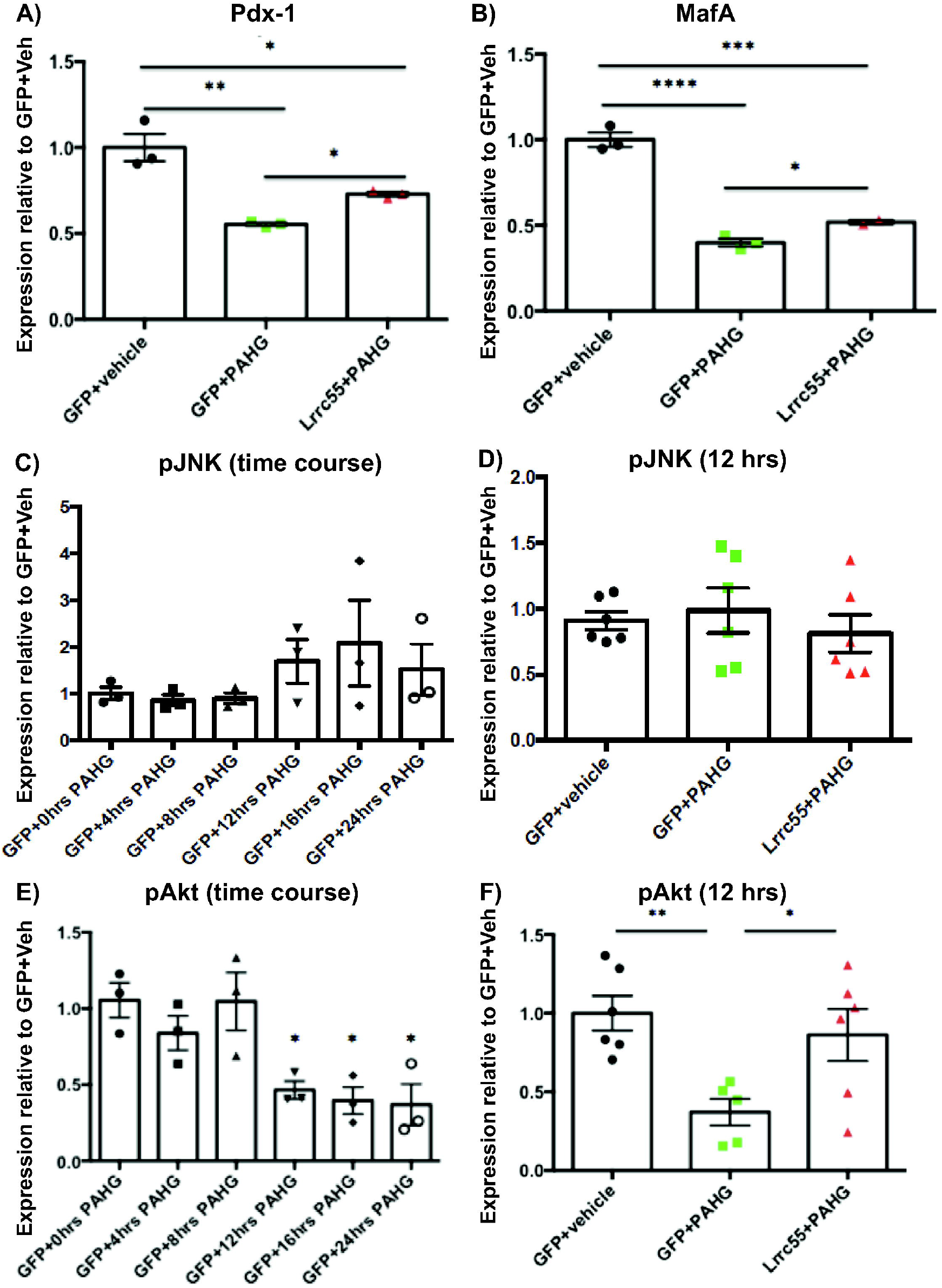
(A, B) Dissociated mouse islets were infected with adenovirus expressing GFP (AdGFP) or Lrrc55 (AdLrrc55) then treated with 0.5 mM palmitate and 33.3 mM glucose (PAHG) for 72 hrs. Pdx-1 (A) or MafA (B) mRNA expression was measured by RT-qPCR and normalized to expression of mouse PGK1 (housekeeping gene). Results are presented as means ± SEM of 3 independent experiments. Each experiment uses islets pooled from 6-7 mice. ** p< 0.01, * p<0.05 (unpaired student’s T-test between two groups). (C-F) INS-1 cells were infected with AdGFP or AdLrrc55 and cultured in 1mM palmitate and 33.3 mM glucose for the indicated time. Protein expression was measured by western immunoblotting and normalized to actin. Results are presented as means ± SEM of 3-6 independent experiments. “**”= p< 0.01, “*”= p<0.05 (unpaired student’s T-test between two groups).

Next, we examined the expression of JNK and Akt. JNK has a role in ER stress signaling, insulin synthesis, and beta-cell function^21^. Akt is a pro-survival gene and its expression is reduced in response to GLT in beta-cells ^22^. Akt downregulation is associated with the upregulation of the BK channel ^23^. Interestingly, using INS-1 cells treated with Ad-GFP and palmitate for 24 hrs, we observed a small increase in phospho-JNK expression after 12 hours of PA exposure, which reverted to baseline at 24 hrs. Overexpression of Lrrc55 had a small but not statistically significant effect on phospho-JNK expression. In contrast, phospho-Akt expression was significantly down regulated in INS-1 cells exposed to palmitate; overexpression of Lrrc55 can rescue pAkt expression after a 12-hr but not a 16-hr palmitate exposure (Figure 1 C-F).

### Lrrc55 partially reversed GLT-induced changes in ER calcium channel expression

Previously, we found that overexpression of Lrrc55 attenuated the GLT-induced depletion of releasable ER calcium stores in beta cells ^6^. Since ER calcium stores are maintained by SERCA, IP3R, and RyR2, we first determined the expression of these transporters. We found that expression of SERCA2a, SERCA2b, and SERCA2c were all downregulated in islets after 72 hours exposure to GLT, which were partially restored by overexpression of Lrrc55 (Figure 2A-C). Interestingly, overexpression of Lrrc55 did not restore GLT-induced reduction in IP3R1 or IP3R3 expression (Figure 2 D,E). Moreover, overexpression of Lrrc55 reversed GLT-induced upregulation of RyR2 expression in islets (Figure 2 F).

**Figure 2.**
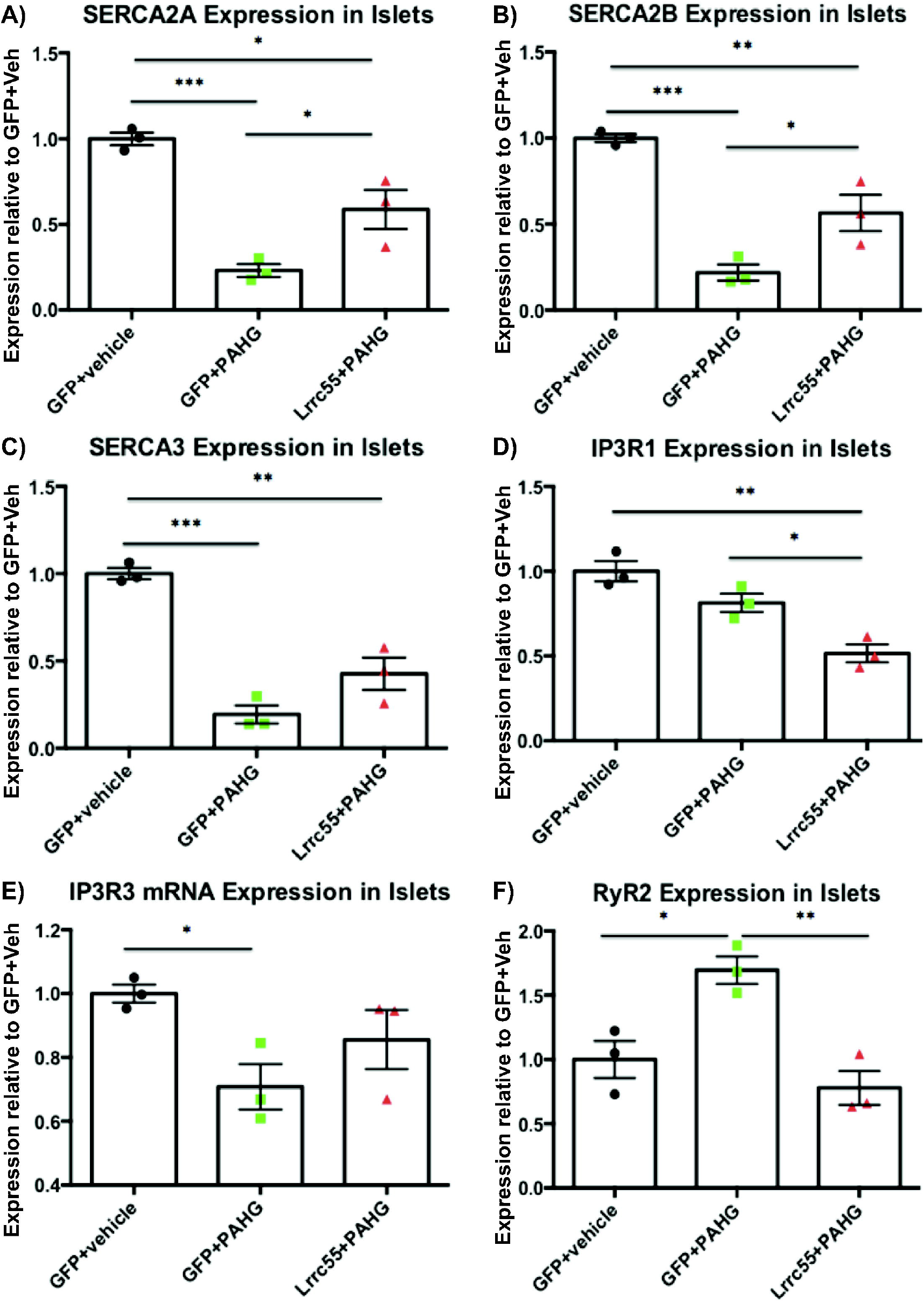
Dissociated mouse islets were infected with AdLrrc55 or AdGFP then treated with 0.5 mM palmitate (in 1% BSA) and 33.3 mM glucose (PAHG) or control (vehicle, 1%BSA) for 72 hours. mRNA expression of SERCA2A (A), SERCA2B (B), SERCA3 (C), IP3R1 (D), IP3R3 (E), RyR2 (F) were measured by RT-qPCR and normalized to PGK1 (housekeeping gene). Results are presented as means ± SEM of 3 pooled independent experiments. Each experiment used islets pooled from 5-6 mice. “**”= p< 0.01, “*”= p<0.05 (unpaired student’s T-test between two groups).

### Lrrc55 on ER calcium channel activity

Next, we examined whether Lrrc55 overexpression modulates activity of SERCA and IP3R. We transfected HEK 293T cells to overexpress Lrrc55 and SERCA2B, chosen because it is highly expressed in beta cells ^14,15^. We found that expression of Lrrc55 does not affect SERCA activity (Figure 3 A, B). We co-expression Lrrc55 and IP3R in Flp-In T-REx 293T cells overexpressing IP3R and again, Lrrc55 does not appear to affect IP3R activity (Figure 3 C, D).

**Figure 3.**
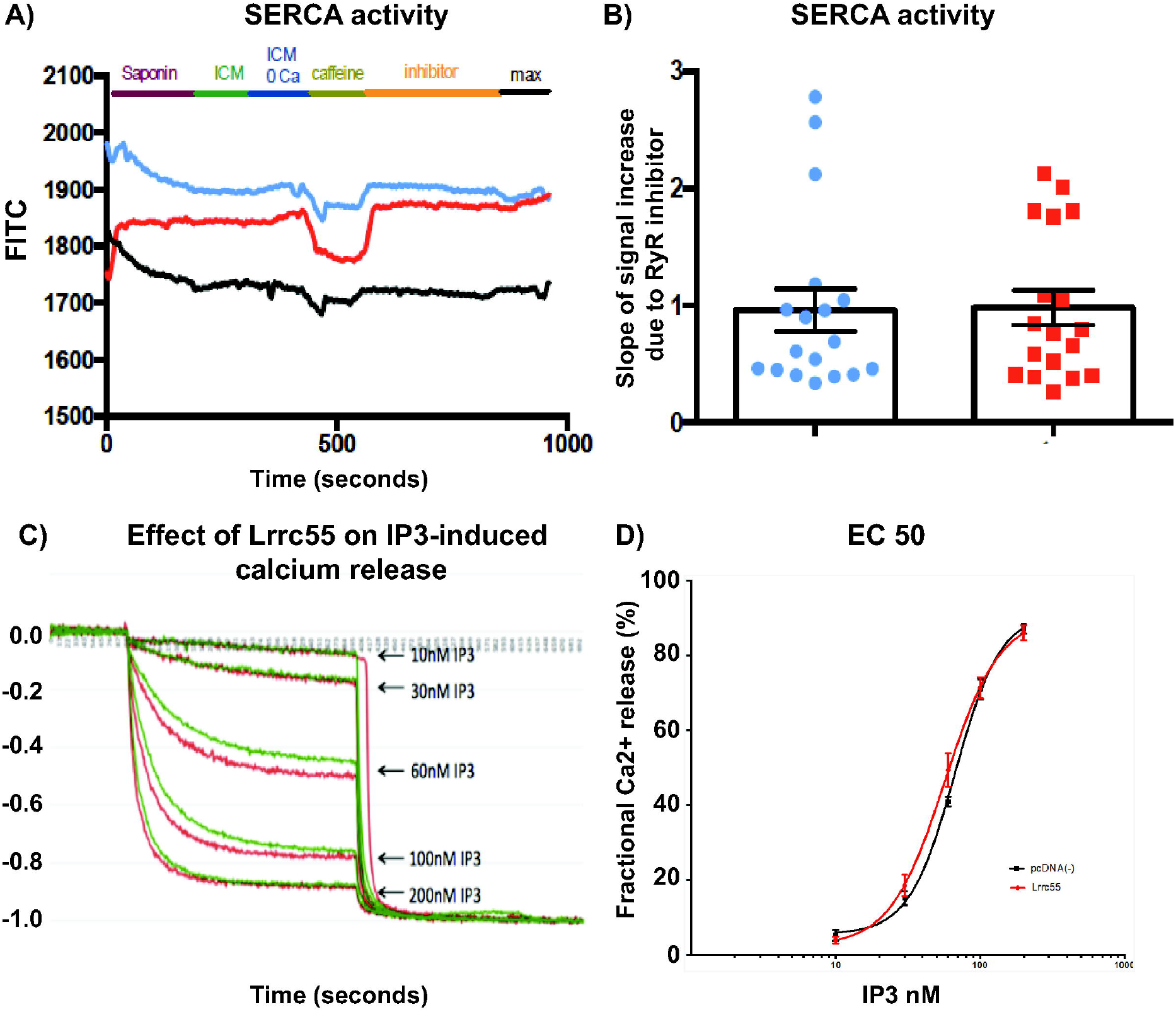
(A, B) HEK 293T cells were transfected with plasmids containing G-CEPIA1er, SERCA2b, RyR2, and Lrrc55 or a negative vector (pcDNA(−/−)), then stimulated with caffeine to release ER calcium, and subsequently RyR was blocked by specific inhibitors to prevent calcium leak from the ER. (A) Representative image of a SERCA activity is presented here. (B) Rate of ER calcium increase due to SERCA activity. Results are presented as means ± SD of 18 coverslips each, with a minimum of 10 cells quantified for each condition, from 5 independent experiments. (C, D) Flp-In T-Rex 293T cells were transfected with plasmids containing G-CEPIA1er, and Lrrc55 or a negative vector (pcDNA(−/−)), then stimulated with IP3 to release ER calcium, ER calcium levels is reflected by G-CEPIA fluorescence. (C) Representative of a IP3R activity is presented. (D) Fractional release of ER calcium resulting from various doses of IP3. Results are presented as means ± SD of 7 independent experiments.

### Lrrc55 on susceptibility to STZ-induced DM

Our published work showed that Lrrc55 is a pro-survival factor, and it is upregulated by >60 fold in islets during pregnancy. Therefore, we examined whether Lrrc55 is protective against streptozotocin-induced beta-cell death and diabetes. We administered streptozotocin to pregnant wild type or Lrrc55KO mice on days 13, and 14 of pregnancy, and measured blood glucose for the following 4 weeks, starting at 1 week after the last dose of streptozotocin. Here, we found that wild type mice are less susceptible to streptozotocin-induced diabetes (Figure 4 A). Interestingly, in mice that did not become diabetic, there was a trend for higher blood glucose during an IPGTT in the Lrrc55 KO mice (Figure 4 B).

**Figure 4.**
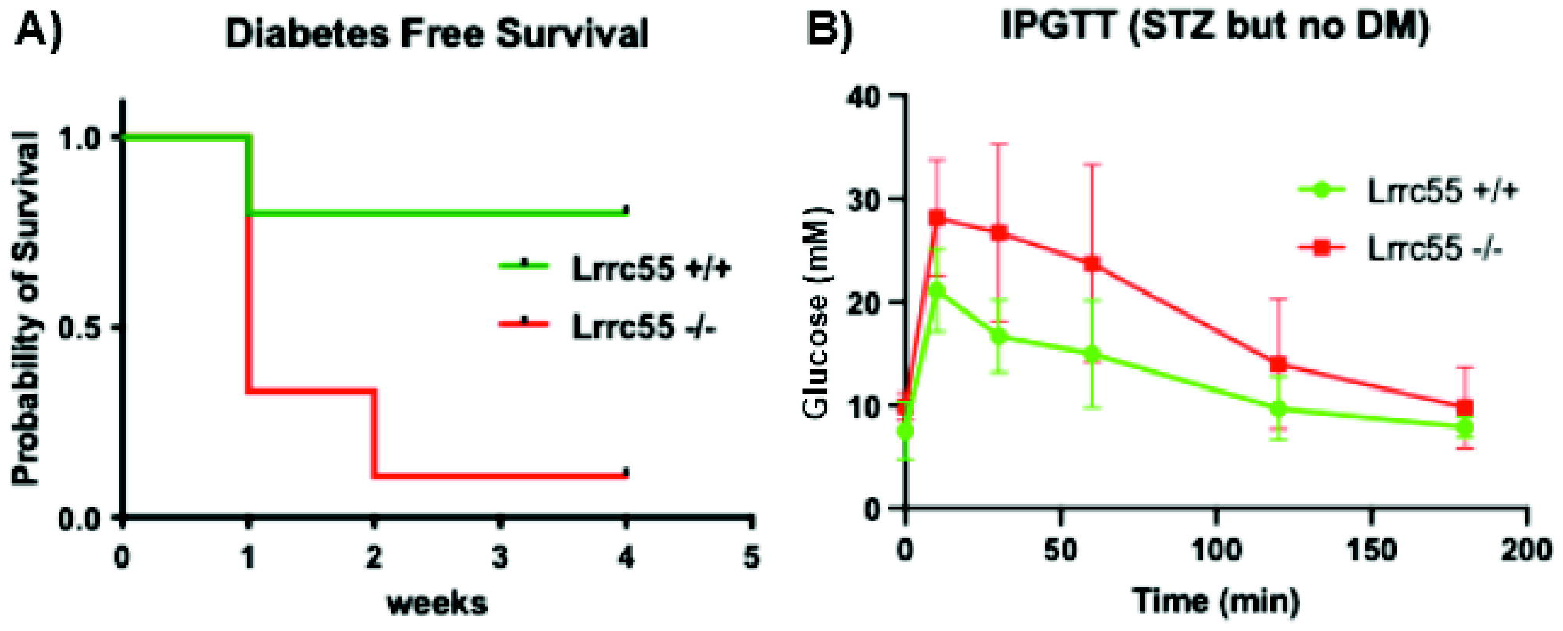
(A) Female pregnant mice were given streptozotocin on day 13 and 14 of pregnancy, and then blood glucose was monitored twice weekly. Blood glucose >11mM for two consecutive weeks defined presence of diabetes. Wild type (Lrrc55^+/+^, n=6 mice), mutant (Lrrc55^−/−^: n=9 mice). P=0.019 (Logrank test). (B) IPGTT in mice given streptozotocin but did not become diabetic. Wild type (Lrrc55^+/+^, n=5 mice), mutant (Lrrc55^−/−^: n=4 mice).

### Lrrc55 and GDM

Interestingly, during pregnancy, the Lrrc55KO mice has better glucose tolerance than the wild type mice (Figure 5 A, B). This is accompanied by a more robust insulin secretory response, both expressed as insulinogenic index and as area-under-the curve throughout the OGTT (Figure 5 C, D).

**Figure 5.**
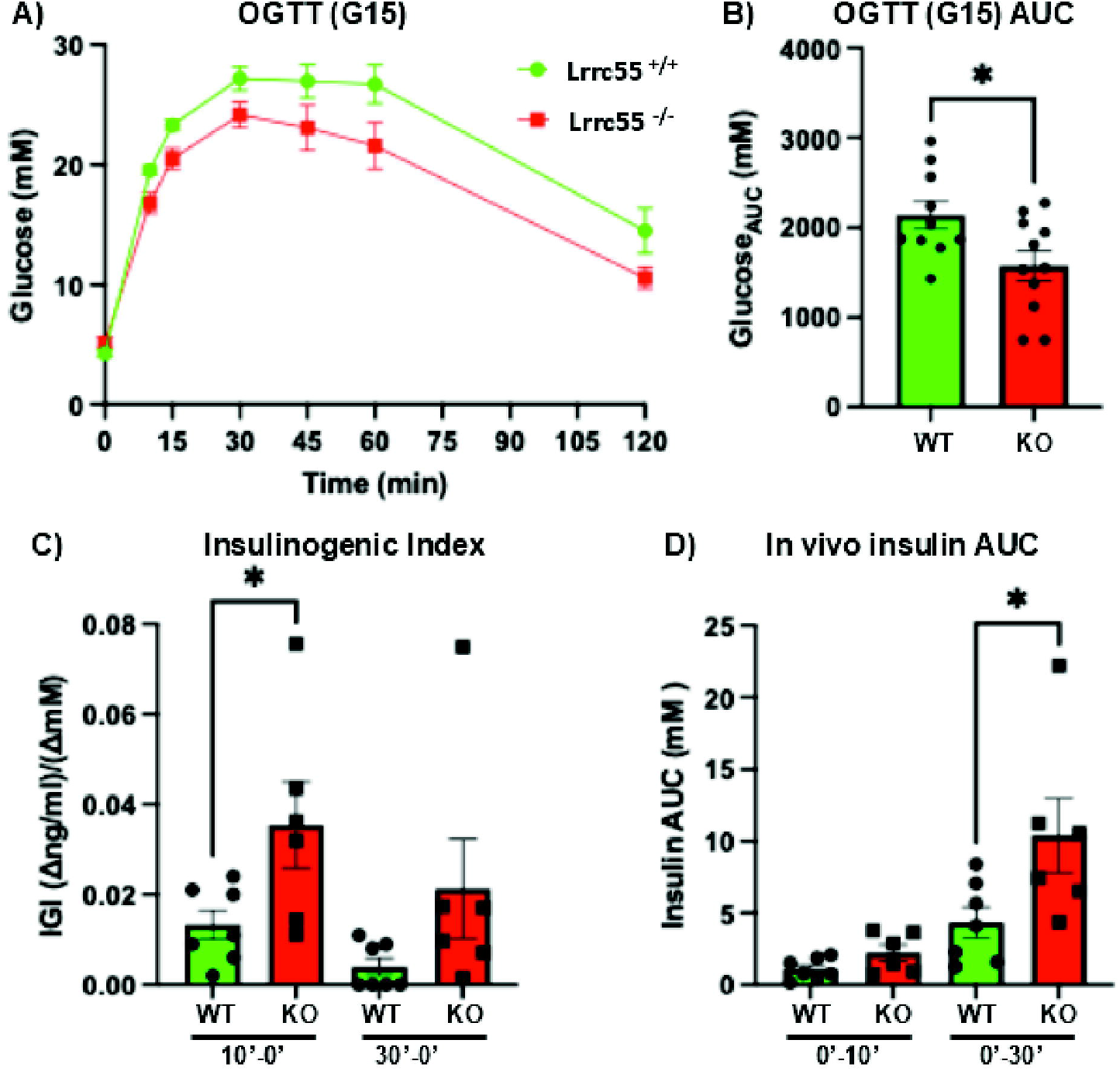
(A,B) Oral glucose tolerance in female wild type (Lrrc55^+/+^) and mutant (Lrrc55^−/−^) mice on day 15 of pregnancy; n=10-11 mice. (C) Insulinogenic index (change in insulin levels relative to change in glucose level, from times 0 to 10 minutes of OGTT [10’-0’], and from time 0 to 30 minutes of OGTT [30’-0’]); n=5-7 mice/genotype. (D) In vivo insulin levels during OGTT, expressed as area under the curve (AUC) from times 0 to 10 minutes of OGTT (0’-10’) and 0 to 30 minutes of OGTT (0’-30’); n=5-7 mice/genotype.

## Discussion

Beta-cell adaptation to the physiologic insulin resistance of pregnancy requires action of prolactin receptor. Previously, we identified Lrrc55 as a novel downstream target of Lrrc55. It is highly expressed in the brain but during pregnancy, its expression is highly and specifically up regulated in the islets. Overexpression of Lrrc55 rescued glucolipotoxicity (GLT)-induced b-cell apoptosis. This was accompanied by an up regulation of prosurvival factors and down regulation of pro-apoptotic genes. Concomitantly, we observed a partial restoration of the glucolipotoxicity-induced reduction in ER calcium store. Here, we demonstrated the indeed, Lrrc55 can reverse some of the GLT-induced changes in calcium channels in b-cells. More interestingly, and unexpectedly, we observed that female Lrrc55^−/−^ mice were more glucose tolerant than their wild type littermates and secreted less insulin during a glucose tolerance test, suggesting that under physiologic conditions, Lrrc55 negatively regulate insulin secretion.

In this study, we explored two potential mechanisms by which Lrrc55 might increase β-cell ER calcium stores in order to mediate survival: by altering the activity of either 1) the Sarco/Endoplasmic Reticulum Ca2+-ATPase (SERCA) or the 2) the Inositol Triphosphate Receptor (IP3R). our results indicated that Lrrc55 did not affect either SERCA or IP3R activity in HEK cells under basal conditions. Another potential mechanism could be to alter the expression levels of these channels. There are multiple isoforms of SERCA, 3 of which are expressed in pancreatic β-cells: SERCA2A, SERCA2B, and SERCA3 ^14,15^. In dissociated mouse islets, we found a significant decrease in expression of all 3 SERCA isoforms. The down regulation of SERCA2A and SERCA2B, but not SERCA3, was partially restored by Lrrc55 expression, providing a potential mechanistic explanation for how Lrrc55 increases ER calcium stores in conditions of GLT, as this would correspond to a higher concentration of calcium pumped from the cytosol into the ER. Expression of two IP3R isoforms were also evaluated: IP3R1 and IP3R3, as these are the isoforms, with IP3R3 being most highly expressed in pancreatic β-cells ^9^. In mouse islets, IP3R1 expression was unchanged by exposure to GLT, but significantly decreased when Lrrc55 is overexpressed. This provides a plausible mechanism of maintaining ER calcium stores, as a lower level of IP3R1 would reduce of the number of channels available for calcium to be released from the ER back into the cytosol. RyR2 is the major isoform of RyR present in β-cells ^8^. Interestingly, in mouse islets, RyR2 RNA expression was increased by GLT, which was attenuated by Lrrc55. This provides a third mechanism by which Lrrc55 can increase ER calcium stores, as a decrease in RyR2 expression may lead to a decrease in calcium release from the ER into the cytosol. Consistent with the putative role of Lrrc55 in preserving ER calcium stores and protect β-cells against GLT-induced apoptosis, we found that pregnant Lrrc55^−/−^ are more susceptible to streptozotocin-induced diabetes. Streptozotocin is a β-cell toxin the gains cell entry exclusively through the glucose transporter, GLUT2, which is abundantly and mainly expressed on β-cells. Moreover, mice that received streptozotocin but never developed diabetes had higher blood glucose, suggest that Lrrc55 preserves β-cell function in the presence of a β-cell toxin. Most interestingly, when we examined glucose tolerance and insulin secretion of pregnant mice, a condition associated with significant up regulation of Lrrc55, we found that presence of Lrrc55 negatively impact glucose tolerance and insulin secretion. Of note, we have previously shown that during pregnancy, we did not detect any apoptotic cells in the pancreatic islets. Lrrc55 is an auxiliary subunit of BK channel, and it causes a large negative shift in voltage-dependent BK channel activation^4^, suggesting it may affect BK channel activity. BK channels regulate membrane repolarization and both stimulatory and inhibitory role in β-cell membrane depolarization has been found^24-26^.

In summary, Lrrc55 appears to regulate ER calcium channel expression in pancreatic islets under condition of glucolipotoxicity. While it does not affect calcium channel activity under basal condition, its effect under stressed conditions is unknown. Most intriguingly, Lrrc55 appears to hinder insulin secretion in vivo during pregnancy. Whether this is due to intrinsic effect on β-cells, or systemic effect in non-β-cells will need further investigation. Lrrc55 is highly expressed in neurons, which affects in vivo insulin secretion. The potential role of Lrrc55 in neurons and its potential secondary effect in β-cell function needs to be considered.

## Acknowledgement

This work was supported by funding from the Natural Sciences and Engineering Research Council of Canada (NSERC) -RGPIN-2020-05247 to Carol Huang. Raneet Kahlon was supported by a NSERC summer studentship and an O’Brien’s summer studentship. We thank Dr. SR Wayne Chen and Dr. Jonathan Lytton for guidance with experimental design and provide technical advice.

